# “Extensive Transmission of Microbes along the Gastrointestinal Tract”

**DOI:** 10.1101/507194

**Authors:** TSB Schmidt, MR Hayward, LP Coelho, SS Li, PI Costea, AY Voigt, J Wirbel, OM Maistrenko, RJ Alves, E Bergsten, C de Beaufort, I Sobhani, A Heintz-Buschart, S Sunagawa, G Zeller, P Wilmes, P Bork

## Abstract

The gastrointestinal tract is abundantly colonized by microbes, yet the translocation of oral species to the intestine is considered a rare aberrant event, and a hallmark of disease. By studying salivary and fecal microbial strain populations of 310 species in 470 individuals from five countries, we found that transmission to, and subsequent colonization of, the large intestine by oral microbes is common and extensive among healthy individuals. We found evidence for a vast majority of oral species to be transferable, with increased levels of transmission in colorectal cancer and rheumatoid arthritis patients and, more generally, for species described as opportunistic pathogens. This establishes the oral cavity as an endogenous reservoir for gut microbial strains, and oral-fecal transmission as an important process that shapes the gastrointestinal microbiome in health and disease.

## Main Text

Both the oral cavity and large intestine accommodate unique microbiomes that are relevant to human health and disease (Lynch and Pedersen, 2016; Wade, 2013). Mouth and gut are linked by a constant flow of ingested food and saliva along the gastrointestinal tract (GIT), yet they host distinct microbial communities (Ding and Schloss, 2014; Segata et al., 2012) in distinct microenvironments (Savage, 1977), and have been reported to harbour locally adapted strains (Lloyd-Price et al., 2017).

The segregation of oral and intestinal communities is thought to be maintained by various mechanisms, such as gastric acidity (Howden and Hunt, 1987; Martinsen et al., 2005) and antimicrobial bile acids in the duodenum (Ridlon et al., 2014). Failure of this oral-gut barrier has been proposed to lead to intestinal infection (Martinsen et al., 2005), and the prolonged usage of proton pump inhibitors can result in an enrichment of particular oral microbes in the gut (Imhann et al., 2016). Increased presence of specific oral taxa in the intestine has in turn been linked to several diseases, including rheumatoid arthritis (Zhang et al., 2015), colorectal cancer (Flynn et al., 2016; Zeller et al., 2014) and inflammatory bowel disease (IBD, (Gevers et al., 2014)). While it remains unclear whether disease-associated strains are indeed acquired endogenously (from the oral cavity) or from the environment, it was recently shown that *Klebsiella* strains originating from salivary samples of two IBD patients triggered intestinal inflammation in gnotobiotic mice (Atarashi et al., 2017).

This suggests that the presence of oral commensals in the gut is a rare, aberrant event as a consequence of *ectopic* colonization (i.e., ‘in the wrong place’), and hence a hallmark of disease. Outside a disease context, however, possible links between the oral and gut microbiome remain poorly characterized. Several genera were shown to be prevalent at both sites (Segata et al., 2012), with community types in one being weakly predictive of the other (Ding and Schloss, 2014), and with similar gene content in particular species (Franzosa et al., 2014), but with distinct, locally adapted strains (Lloyd-Price et al., 2017). We hypothesised that this picture is incomplete, and that microbial transmission along the GIT is more common than previously appreciated: that despite oral-gut barrier effects, some microbes freely and frequently traverse the GIT and colonise different niches, forming continuous populations that shape the human microbiome.

To test this hypothesis, we assembled and analyzed a dataset of 753 public and 182 newly sequenced saliva and stool metagenomes from 470 healthy and diseased individuals (diagnosed with rheumatoid arthritis, colorectal cancer or type-1 diabetes) from Fiji (Brito et al., 2016), China (Zhang et al., 2015), Luxembourg (Heintz-Buschart et al., 2016), France (Zeller et al., 2014), and Germany (Voigt et al., 2015) (see Methods, Figure S1, and table S1). For these samples we profiled 310 prevalent species, accounting for 99% of classifiable microbial abundance in both saliva and stool (see Methods and table S2). We reasoned that if transmission between the oral and gut microenvironments is frequent, we would expect salivary and fecal microbial populations to be more similar within an individual than between individuals. Conversely, under a strong barrier model with restricted transmission, intra- and inter-individual similarities would be equivalent.

We found that at species level, community composition was consistent with distinct populations occupying the oral and intestinal microenvironments. By prevalence across subjects, the 310 profiled species fell into three categories (Figure 1A): 44% were predominantly fecal (observed in ≥10% of fecal, but <10% of saliva samples), including core members of the gut microbiome, such as *Clostridium sp., Ruminococcus sp.* and *Bacteroides sp*.; 16% of species were predominantly oral. Although the remaining 125 (40%) species were prevalent in ≥10% of saliva and stool samples, their relative abundances differed greatly between the two habitats. The overall oral and fecal microbiome compositions appeared independent of each other (between-subject Bray-Curtis dissimilarities per site, ρ_Pearson_=-0.03), and the compositional overlap between mouth and gut of the same subject was not found to be significantly different when compared to a between-subject background (Wilcoxon test, Bray-Curtis dissimilarities, p=0.46).

**Figure 1:**
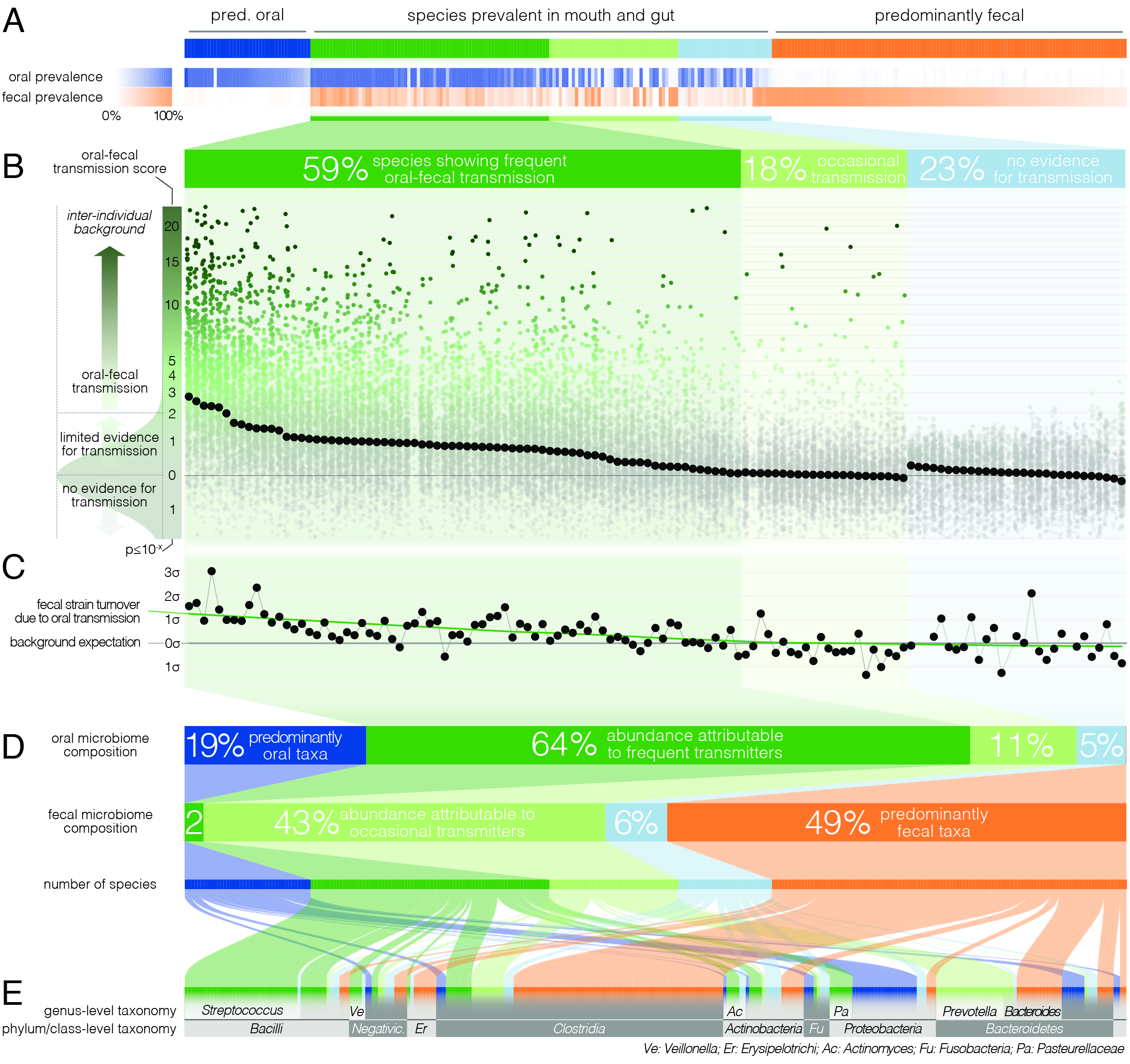
Oral-fecal transmission is common across a wide range of phylogenetically diverse species. (A) Among 310 tested species, 125 were prevalent in both the mouth and gut across subjects. (B) 77% of these formed coherent strain populations between both habitats, when viewed across all tested subjects (‘frequent’ transmitters) or at least in some (‘occasional’ transmitters), as evidenced by oral-fecal transmission scores based on intra-individual SNV overlap against an inter-individual background (see Methods). (C) Oral-to-fecal transmission rates, as inferred from longitudinal coupling of oral and gut SNVs (see Methods), exceeded background levels for transmitted taxa, even at conservative lower estimates. (D) On average, transmissible taxa accounted for a large fraction of classifiable microbial abundance in both the oral cavity and gut. (E) Oral-fecal transmissibility was largely a clade-wise trait at genus or family ranks, but common across bacterial phyla.

However, to accurately establish and quantify microbial transmission, it is necessary to track populations at the resolution of strains rather than species, as demonstrated previously in fecal microbiota transplantation (Li et al., 2016) or seeding of the infant microbiome (Asnicar et al., 2017; Korpela et al., 2018). We therefore profiled microbial single nucleotide variants (SNVs) across metagenomes, as a proxy for strain populations (Li et al., 2016). We formulated a transmission score for each species per subject, based on the likelihood that the observed intra-individual SNV overlap was generated by an inter-individual background model (see Methods). Of the 125 species prevalent in both mouth and gut, 77% showed evidence of oral-fecal transmission. Out of these, 74 species (59%) showed significantly higher intra-individual SNV similarity across all subjects compared to cohort-wide background SNV frequencies (Benjamini-Hochberg-corrected Wilcoxon tests on transmission scores, p<0.05, see Methods; Figure 1B, S2, table S2). This suggests that they form coherent strain populations along the GIT in most subjects, subject to frequent oral-fecal microbial transmission. Strains of *Streptococcus, Veillonella, Actinomyces* and *Haemophilus*, among other core oral taxa, fell into this category. An additional 22 species (18%) showed evidence of at least occasional transmission, with individually significant oral-fecal SNV overlap in some, but not across all subjects, as did 18 species that were generally prevalent in either the mouth or the gut (but not both). All 21 members of the *Prevotella* genus, an important clade of the gut microbiome, were among these occasionally transmitted species. The remaining 29 (23%) species, which were prevalent in both sites, did not show signs of transmission under the strict thresholds we applied.

The fecal abundance of all species with paired observations exceeded lower-bound physiologically predicted levels (i.e., the detection of salivary bacteria in stool purely as the result of ingestion) by several orders of magnitude, even with conservative estimates (Figure S3). An average person swallows an estimated 1.5 * 10^12^ oral bacteria per day (Humphrey and Williamson, 2001; Sender et al., 2016). Passage through the stomach reduces the viable bacterial load by 5-6 orders of magnitude (Giannella et al., 1972; Sender et al., 2016), a reduction that is expected to be mirrored at the DNA level, given that free DNA, released from dead bacterial cells, is degraded within seconds to minutes in saliva, the stomach and the intestine (see e.g. Mercer et al., 1999 and Liu et al., 2015). Relative to the ~3.8*10^13^ bacterial cells in the large intestine, ‘passive’ transmission without subsequent colonization in the gut would therefore account for a reduction in relative abundance by ~4*10^−7^ from saliva to feces (Figure S1C). Thus, the observed overlap of microbial SNVs could not be explained by passive translocation, but was indeed caused by active colonization in the gut. Moreover, transmission scores were independent of technical covariates, such as the horizontal or vertical coverage of genome mappings (Fig S4). Average transmission scores per species across subjects correlated with prevalence in saliva (ρ_Spearman_=0.6) but not stool (p=0.05), reinforcing the notion that core oral taxa tended to be transmitted. Given the limited microbial read depth of salivary metagenomes (due to high fractions of human DNA), this result also indicates that our estimates of oral-fecal transmissibility were quite conservative, with potentially high rates of false negatives.

It was recently shown that during early life, infants are colonised by maternal strains from both the oral cavity and gut (Ferretti et al., 2018), and that strains from the latter can persist in the infant gut at least into childhood (Korpela et al., 2018). Therefore, to determine whether the observed intra-individual overlap of selected strain populations was due to continuous oral-gut transmission or rare colonization events with subsequent independent expansion in each site, we focused on a subset of 46 individuals for whom longitudinal data was available (with sampling intervals ranging from 1 week to >1 year; mean 79 days). We found that both oral and fecal strain populations were usually stable, even over extended periods of time (Fig S5), in line with earlier observations for each individual body site (Lloyd-Price et al., 2017; Schloissnig et al., 2012). Oral and fecal longitudinal SNV patterns were coupled for transmitted species (see Methods): oral SNVs observed at an initial time point were significantly enriched among fecal SNVs that were newly gained over time, but generally not vice versa (Fig S6). Moreover, oral-fecal transmission rates (i.e., the fraction of fecal strain turnover attributable to oral strains; see Methods) significantly exceeded background expectation for frequently transmitted taxa (Fig 1C). These findings orthogonally support the oral-gut transmission hypothesis as they strongly suggest that transmission is in the direction of mouth to gut, and not vice versa; and they imply that oral-intestinal transmission is indeed a frequent and continuous process in which oral strain populations constantly re-colonize the gut.

Oral-fecal transmissibility, as a trait, generally aligned with phylogenetic clade boundaries (phylogenetic signal, λ_Pagel_=0.76), although transmitting groups were found across bacterial phyla (Fig 1DE, S3, table S2). Transmission scores were negatively correlated with genome size (ρ_Spearman_=−0.6), indicating that transmitted species generally had smaller genomes than non-transmitted ones. Moreover, oxygen tolerant species (aerobes and facultative anaerobes) showed 7-fold higher scores than anaerobes on average (ANOVA, p=10^−16^). In contrast, no association was observed for sporulation and motility. To account for possible bias in the species reference and the phylogenetic signal of oral-fecal transmissibility, we confirmed that these signals were robust to phylogenetic regression (table S2).

Viewed across individuals, we found that seeding of the gut microbiome from the oral cavity was extensive, with high levels of variation (Fig 2A). On average, potentially transmissible species (i.e., frequent and occasional transmitters) accounted for 75% of classifiable microbes in saliva, ranging up to 99% in some subjects. However, not all of these were detectable in the matched fecal samples, and oral-fecal strain overlap was generally incomplete. We therefore quantified the fraction of *realised* transmission based on paired observations of species and intra-individual SNV overlap (see Methods). With these criteria, on average 35% of classifiable salivary microbes were transmitted strains that could be traced from mouth to gut within subjects. Similarly, on average 45% (range 2%-95%) of classifiable fecal microbes were potential transmitters. These included common fecal species (e.g., *Prevotella copri*) that were detectable in a subset of salivary samples and showed only occasional transmission. Nevertheless, on average only 2% of classifiable fecal microbes could be confidently ascribed to transmitted strains, ranging to >30% in some subjects.

**Figure 2:**
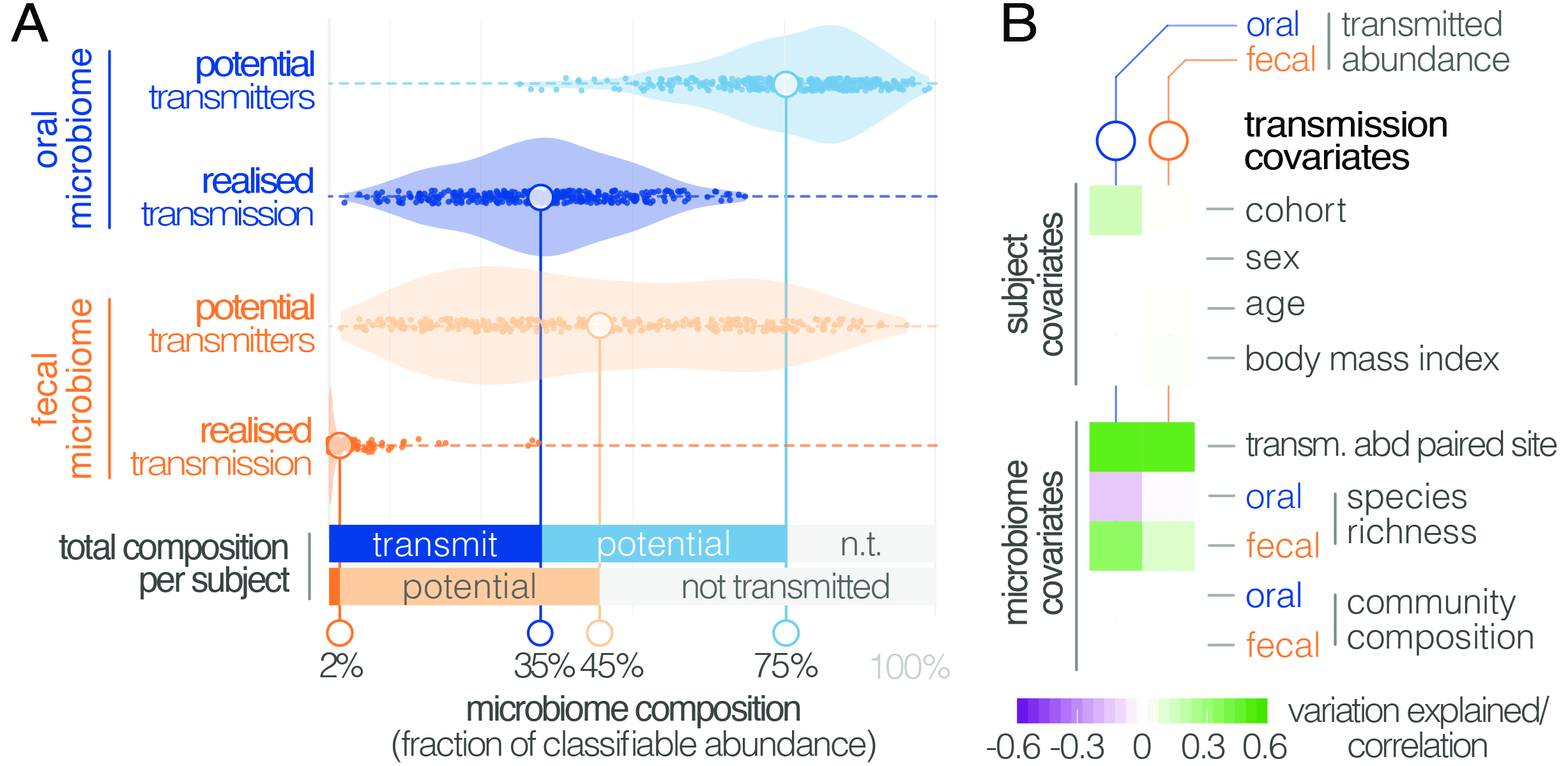
Oral-fecal transmission is extensive, with high levels of variation across individuals. (A) Potentially transmissible species on average accounted for 75% and 45% of known microbes in salivary and fecal samples, respectively. Among these, realised transmitters were defined as strains that could be traced within subjects with confidence (given detection limits, see Methods). (B) Tests for the association of transmission levels in mouth and gut to subject-level covariates (ANOVA, relative sum of squares), to each other (ρ_Spearman_), with oral and fecal community richness (ρ_Spearman_), and with oral and fecal community composition (distance-based redundancy analysis on Bray-Curtis dissimilarities, blocked by cohort, relative sum of squares).

Between-subject variation in the relative abundance of transmitted oral and fecal microbes was found to be independent of subject sex, age and body mass index, although moderate differences were observed between study cohorts (ANOVA, p=0.002; Figure 2B; table S3). Levels of transmitted microbial abundance in mouth and gut were found to correlate with each other (ρ_Spearman_=0.48) and with fecal species richness, but salivary transmitted abundance negatively correlated with oral species richness. This is in line with the observation that core oral species are transmissible, with higher richness implying the increased presence of non-transmitted taxa. Conversely, transmission would add species to a mostly non-transmissible core community in the gut.

Although there was no overall association to community composition, levels of transmission correlated with oral or fecal abundances of individual genera (table S3). To test whether specific oral and gut microbiome features were predictive of transmission, we categorized individuals based on total transmitted abundance in saliva and stool as ‘high’ or ‘low’ transmission individuals (Methods). We found that models based on salivary species abundances were mildly predictive of both oral (AUC=0.738) and fecal (AUC=0.642) transmission levels (table S4, Figure S7). Gut species models, in contrast, were very strong predictors of transmission in both mouth (AUC=0.951) and gut (AUC=0.971). This signal was largely driven by the enrichment of transmitting species in stool (table S4), but surprisingly robust to an elimination of all detected transmitters from the model (AUC=0.835 for the stool transmission group), again implying that the true extent of oral-intestinal transmission may indeed exceed our conservative estimates. *Fusobacterium nucleatum subsp. animalis* and *nucleatum* stood out among non-trivial gut markers enriched in high-transmission individuals, in line with existing hypotheses that *Fusobacterium nucleatum* subspecies may enable synergistic colonization of oral bacteria in the gut, in association with certain diseases (see e.g. Flynn et al., 2016).

In general, the fecal enrichment of specific oral microbes has repeatedly been associated with various diseases (Zeller et al., 2014; Zhang et al., 2015). However, due to insufficient taxonomic resolution, oral provenance has so far remained impossible to distinguish from an influx of closely related but distinct strains from the environment. We therefore defined a list of disease states with putative links to oral-fecal transmission and annotated known associations in the literature to all species in our dataset (Fig 3A; table S2). Transmission scores were significantly increased for known opportunistic pathogens (ANOVA, p=0.016), causative agents of dental caries (p=10^−9^), and plaque-dwelling bacteria (p=0.002). Likewise, species associated with periodontitis showed increased evidence for transmission (p=0.002), though this signal was mostly due to mildly periodontic species, while core drivers, such as *Tannerella forsythia, Treponema denticola* and *Porphyromonas gingivalis* (Socransky et al., 1998), showed little or no indication of oral-fecal transmission. Endocarditis-associated species showed significantly increased transmission scores upon phylogenetic regression (p=0.007), mostly driven by *Haemophilus, Aggregatibacter* and viridans Streptococci. This overall elevated transmissibility of taxa known to colonize ectopically in various habitats across the body (i.e., opportunistic pathogens), in particular via the bloodstream and associated with inflammation (i.e., endocarditis- or periodontitis-associated species (Hajishengallis, 2014)), may provide first cues to possible mechanisms of oral-fecal transmission.

**Figure 3:**
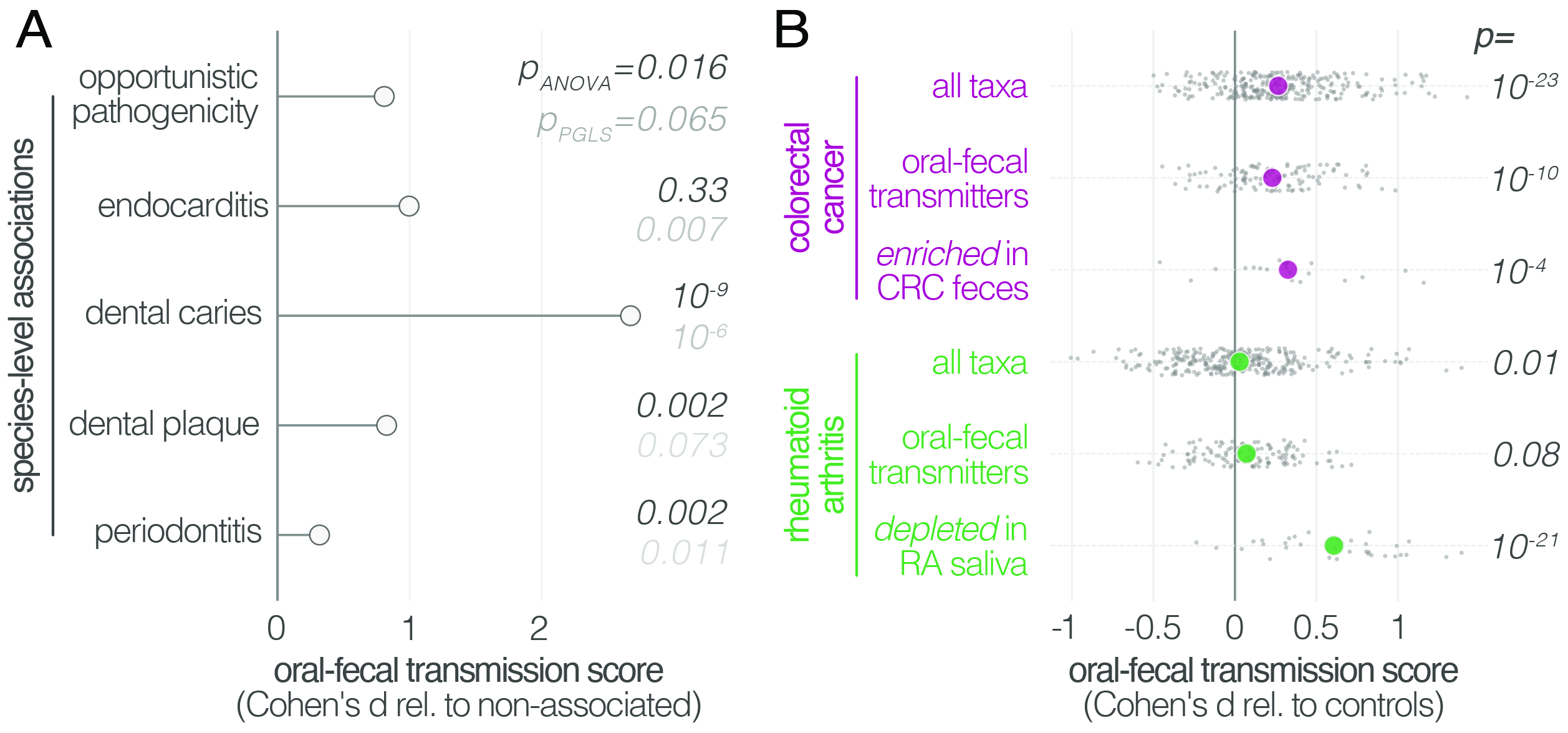
Oral-fecal transmission is associated with disease state. (A) Species known to be associated with various diseases showed increased oral-fecal transmission scores (p_anova_, sequential ANOVA including additional phenotypes), even upon phylogenetic generalized least squares regression (p_pgls_, see Methods and table S2). (B) Oral-fecal transmission scores tested in colorectal cancer and rheumatoid arthritis cases against controls for specific sets of species (sequential ANOVA, blocked by taxon and subject covariates). Individual data points represent Cohen’s d effect sizes (difference in means, normalised by pooled standard deviation) for individual taxa across subjects.

Our dataset included metagenomes from case-control studies for rheumatoid arthritis (RA, (Zhang et al., 2015)), colorectal cancer (CRC, (Zeller et al., 2014)) and type-1 diabetes (T1D, (Heintz-Buschart et al., 2016)), totaling 299 individuals, including 172 with salivary and fecal samples. Treatment-naïve CRC patients, sampled before colonoscopy, showed increased transmission scores across all taxa (average per-taxon Cohen’s d=0.27; ANOVA p=10^−23^; Fig 3B), as well as for transmitted taxa only (d=0.23; p=10^−10^). The effect was even more pronounced for species previously described (Zeller et al., 2014) to be enriched in the feces of CRC patients (d=0.33; p=10^−4^; Figure S8), including *Fusobacterium nucleatum* spp., *Parvimonas micra* and *Peptostreptococcus stomatis*. These findings are in line with a recent report that the oral and fecal microbiome are linked in the context of CRC (Felmer et al., 2018), and support the hypothesis (Flynn et al., 2016) that CRC-associated species are sourced intra-individually from the oral cavity.

Treatment-naïve RA patients displayed mildly elevated transmissibility across all taxa (d=0.03, p=0.01) and transmissible taxa only (d=0.07, p=0.08). Interestingly, species that were orally depleted in RA patients showed markedly increased transmission scores (d=0.61; p=10^−21^). In contrast, a trend towards decreased transmission in T1D patients was not statistically significant.

Our results demonstrate that influx of oral strains from phylogenetically diverse microbial taxa into the gut microbiome is extensive in healthy individuals, with a high degree of variation between subjects. We showed that the vast majority of species prevalent in both the oral cavity and gut form connected strain populations along the gastrointestinal tract. Furthermore, by leveraging longitudinal data, we established that transmission from the mouth to the gut is a constant process. Approximately one in three classifiable salivary microbial cells colonize in the gut, accounting for at least 2% of the classifiable microbial abundance in feces. This puts oral-fecal transmission well in the range of other factors that determine human gut microbiome composition (Schmidt et al., 2018). Moreover, we note that by using saliva and feces as metagenomic readouts, we may underestimate colonization by oral microbes of the mucosa, given that fecal microbiome composition is not fully representative of the gastrointestinal tract (see e.g. Zmora et al., 2018). Therefore, and considering that our estimates of both the number of transmissible species and of the fraction of transmissible microbial abundance are conservative lower bounds due to strict thresholding and current detection limits of metagenomic sequencing, we posit that true levels of transmission are likely even higher, and that virtually all known oral species can translocate to the intestine at least under some circumstances.

Finally, we found increased transmission linked to some diseases, and showed for colorectal cancer and rheumatoid arthritis that disease-associated strains of several species enriched in the intestine are indeed sourced endogenously, i.e. from the patient’s oral cavity, and not from the environment. These results may extend to other diseases beyond those tested here, calling for revised models of microbiome-disease associations that consider the gastrointestinal microbiome as a whole rather than a sum of parts, with important implications for disease prevention, diagnosis, and (microbiome-modulating or -modulated) therapy.

While our findings are observational and do not reveal oral-intestinal transmission routes or mechanistic insights, they challenge current ecological and physiological models of the gastrointestinal tract that assume the oral cavity and large intestine to harbour mostly independent and segregated microbial communities. Instead, most strain populations appear to be continuous along the gastrointestinal tract, originating from the oral cavity, an underappreciated reservoir for the gut microbiome in health and disease.

## Supplementary Figures

**Figure S1: Data and workflow overview**. (A) Oral-fecal transmission scores were calculated from salivary and fecal microbial SNV profiles. (B) Cohort and dataset overview. For longitudinal cohorts (DE-CTR, CN-RA and LU-T1D), both the total number of samples and the number of individuals are shown, as well as the number of individuals considered in time-series analyses. (C) Salivary and fecal microbial loads allow the calculation of physiologically expected levels of "passive” microbial transmission (i.e., by ingestion, without growth). (D) The longitudinal coupling of microbial SNVs between salivary and fecal samples was used to infer transmission directionality and oral-fecal transmission rates (see Methods).

**Figure S2: Phylogenetic distribution of oral-fecal transmission**. A maximum likelihood phylogenetic tree of the species tested in this study (see table S2 and Methods). Annotations, from inside to outside (colour scales as in the main text): fecal species prevalence (fraction of individuals in which the species was detectable in feces); oral prevalence; average transmission score across subjects (see Figure 1C); transmitter category (see Figure 1); fraction of individuals in which the observed transmission score exceeded median background transmission scores. The visualisation was generated using iTol (Letunic and Bork, 2016). Scalable, interactive versions of the full tree and per-phylum subtrees are available online (http://itol.embl.de/shared_projects.cgi; password-less login ‘tsbschm’).

**Figure S3: Enrichment of oral species in the gut**. Relative to physiologically expected levels of ‘passive’ transmission (see Figure S1C), all tested species with paired observations (in saliva and stool of the same individual) showed a fecal enrichment by several orders of magnitude. The fecal enrichment (x axis) is shown on a log2 scale, so values approximate the effective number of cell divisions (without cell deaths) necessary to account for observed fecal levels based on matched oral samples. The left plot shows enrichment purely based on relative salivary and fecal abundance. On the right, the average oral and fecal depths of uniquely mapping reads is used as a reference, normalised by genome size.

**Figure S4: Oral-fecal transmission scores are independent of technical covariates**. Spearman correlations of oral-fecal transmission scores (see Methods) with putative technical covariates per taxon across subjects. Horizontal (breadth) and vertical (depth) genome mapping coverage did not correlate with transmission scores for transmitting taxa, and were anti-correlated for non-transmitting taxa (i.e., deeper coverage reinforced the negative signal for these taxa). In line with this, the total number of observed SNV positions in each site anti-correlated with transmission scores for non-transmitters, and mildly correlated for transmitters. Taxon relative abundance of transmitters in stool tended to correlate positively with transmission scores; arguably, this is a biological rather than a technical effect, as higher transmission rates coincide with higher fecal abundance of transmitted taxa. The same applies to intra-individually shared genome coverage which is likewise expected to coincide with oral-fecal strain overlap.

**Figure S5: Longitudinal stability of SNV profiles per species in saliva and stool**. SNV overlap per taxon and intra-individual time series, normalised as a standard Z score across an inter-individual background. Median Z scores are highlighted.

**Figure S6: Directionality of transmission, as inferred from longitudinal data**. The longitudinal coupling of oral and fecal SNVs was assessed from longitudinal source-sink sample triplets (see Methods, Figure S1). The heatmaps show data on oral-to-fecal (left, blue) and fecal-to-oral (right, orange) coupling. Taxa (y axis) are sorted by transmission category analogous to figure 1 (top to bottom, frequent transmitters, occasional transmitters, non-transmitters, predominantly oral, predominantly fecal); subjects (x axis) are sorted left to right by decreasing evidence for oral-fecal coupling. Colors indicate (significant) positive odds ratios for oral-to-fecal (blue) and fecal-to-oral (orange) coupling, negative odds ratios (grey), or missing/insufficient data (white). Frequent transmitters generally showed indications of oral-to-fecal coupling, but not vice versa. For the remaining taxa, the trend was similar, but less pronounced.

**Figure S7: Multivariable statistical models reveal links between both oral and gut microbiome features with transmission levels**. Models were trained from oral and gut microbiome features to classify subjects into ‘high’ and ‘low’ transmission individuals (see Methods). Model interpretation plots show the median relative model weight (barplots on the left) of the top selected features, the robustness (the number of cross-validation folds in which the respective feature had a non-zero weight; percentages next to the barplot), and the feature z-scores across samples, ordered by group and classification score (heatmap and annotations below). Plots are shown for models trained on the salivary microbiome, predicting the saliva transmission group (a) and the stool transmission group (b); trained on the stool microbiome, predicting the saliva transmission group (c) and the stool transmission group (d); and trained on the stool microbiome after exclusion of frequently transmitting species, predicting the saliva transmission group (e) and the stool transmission group (f). (g) Receiver operating characteristics (ROC) curves for the three models shown in (a, c, e). (h) ROC curves for the three models shown in (b, d, f).

**Figure S8: Species enriched in colorectal cancer show higher oral-fecal transmission scores in patients than controls**. Transmission scores in cases and controls are shown for a list of species previously (Zeller et al., 2014) reported to be fecally enriched in colorectal cancer.

**Figure S9: Horizontal (breadth) and vertical (depth) coverage cutoffs**. To be considered in our study, a species had to meet three criteria in at least 10% of all considered samples: relative abundance >10^−6^; average vertical genome coverage (depth) ≥0.25x; horizontal genome coverage (breadth) ≥5%. The panels show the number of taxa meeting the (A) depth and (B) breadth criterion alone, as a function of coverage. The chosen cutoffs and final number of taxa considered (310) are indicated.

## Supplementary Tables

**Table S1: Sample and subject metadata**. For a subset of individuals in the CN-RA and DE-CTR cohorts, replicates were merged for salivary samples.

**Table S2: Taxa data**. Taxa metadata, annotated disease associations, and raw data on relative abundances, horizontal and vertical coverage of each taxon across all samples.

**Table S3: Transmission covariates**.

**Table S4: Abundances of oral and fecal marker species are predictive of transmission levels**.

## Methods

### Metagenomic Datasets

Publicly available raw sequence data was downloaded from the European Nucleotide Archive (ENA) for the FJ-CTR (FijiCOMP, project accession PRJNA217052) (Brito et al., 2016) and CN-RA (PRJEB6997) (Zhang et al., 2015) cohorts. Sample metadata was parsed from ENA and the respective study publications.

For the LU-T1D (PRJNA289586) (Heintz-Buschart et al., 2016) cohort, newly generated salivary and fecal metagenomes were added under the existing project accession. For the FR-CRC (ERP005534) (Zeller et al., 2014) and DE-CTR (ERP009422) (Voigt et al., 2015) cohorts, newly generated metagenomes were uploaded under project accession PRJEB28422 (samples ERS2692266-ERS2692323).

### Sample Collection

#### German healthy controls (DE-CTR)

Salivary samples were collected at home before dental hygiene and breakfast in the early morning. Donors collected 2-3 ml of saliva and immediately mixed with 15 ml of RNAlater (Sigma-Aldrich). Samples were transported to the laboratory on ice or dry ice and stored at −80C until further processing.

#### French colorectal cancer cohort (FR-CRC)

Subject recruitment and cohort characteristics were described previously (Zeller et al., 2014). Saliva samples were collected in 1.5 ml saline and stored at −80C until further processing.

#### Luxembourg type-1 diabetes cohort (LU-T1D)

Donors collected 2-3 ml of saliva at home before dental hygiene and breakfast in the early morning. Samples were immediately frozen on dry ice, transported to the laboratory and stored at −80C until further processing.

### DNA extraction

#### DE-CTR & FR-CRC

After thawing on ice, 1-2 ml of each sample were centrifuged directly (FR-CRC) or after dilution in RNALater (DE-CTR). Cell pellets were washed 3x in sterile Dulbecco’s PBS (PAA Laboratories) and DNA was extracted using the using the GNOME DNA Isolation Kit (MP Biomedicals). Briefly, cell pellets were lysed using a multi-step process of chemical cell lysis/denaturation, bead-beating and enzymatic digestion as described previously (Zeller et al., 2014). DE-CTR samples were processed in duplicates, with one replicate being enriched for microbial DNA using the NEBNext® Microbiome DNA Enrichment Kit (NEB, Ipswich, USA) following the manufacturer’s instructions.

#### LU-T1D

After thawing on ice, two 500 μl aliquots of each sample were centrifuged. Cell pellets were frozen in liquid nitrogen and lysed by cryo-milling and chemical lysis in RLT buffer (QIAGEN). Cell debris was passed through QiaShredder columns (QIAGEN), before DNA was isolated using the QIAGEN AllPrep kit according to the manufacturer’s instructions, as described previously (Heintz-Buschart et al., 2016).

### Metagenomic Sequencing

Libraries for salivary samples of the French and German cohorts were prepared using the NEBNext Ultra DNA Library Prep kit (New England Biolabs, Ipswich) using a dual barcoding system, and sequenced at 125bp paired-end on an Illumina HiSeq 2000. For the additional LU-T1D samples, libraries were likewise prepared using a dual barcoding system, and sequenced at 150bp paired-end on Illumina HiSeq 4000 and Illumina NextSeq 500 machines.

### Metagenomic Sequence Processing

Raw reads were quality trimmed and filtered against the human genome issue 19 to exclude host sequences using MOCAT2, as described previously (Kultima et al., 2016). For taxonomic profiling, reads were mapped against a database of 10 universal marker genes for 1,753 species-level genome clusters (*specl clusters*, (Mende et al., 2013)), using NGless (Coelho et al., 2018). A maximum likelihood-approximate phylogenetic tree (with the JTT model, (Jones et al., 1992)) for representative genomes of the same 1,753 clusters was inferred based on protein sequences of 40 near-universal marker genes (Mende et al., 2013) using the ETE3 toolkit (Huerta-Cepas et al., 2016), with default parameters for ClustalOmega (Sievers et al., 2011) and FastTree2 (Price et al., 2010).

Metagenomic reads were mapped at 97% sequence identity (across at least 45nt) against full cluster-representative genomes, using the Burrows-Wheeler Aligner *(Li and Durbin, 2009)*, as implemented in NGless (see Supplemental File 1). Reads mapping to multiple genomes at ≥97% identity were discarded from the analysis. Average vertical coverage (sequencing depth) and horizontal coverage (breadth) per microbial genome in each sample were quantified using the qaCompute utility in metaSNV (Costea et al., 2017).

Two cohorts (CN-RA (Zhang et al., 2015) and DE-CTR (Voigt et al., 2015)) contained technical replicates for several salivary samples; these were pooled after the read mapping step.

### Taxa Filtering and Annotation

The dataset was filtered to include taxa satisfying the following criteria in ≥10% of samples (see Figure S9 for details): horizontal coverage (breadth) of ≥0.05; average vertical coverage (depth) ≥0.25; specl cluster relative abundance of ≥10^−6^. These criteria excluded taxa representing 0.8±1.2% of gut and 1.2±1.9% of oral total mapped abundance. For the remaining 310 taxa, general phenotypes (Gram stain, sporulation, motility, oxygen requirement, among others) were annotated using the *PATRIC* database (accessed Dec 2015) (Wattam et al., 2017), and missing values were amended manually. Host and disease association phenotypes (including opportunistic pathogenicity and periodontitis association) were annotated manually, based on published literature and the *MicrobeWiki* website (https://microbewiki.kenyon.edu/index.php/MicrobeWiki, accessed June 2017). Per taxon summary statistics and annotated metadata are available from table S2.

### Identification of Microbial Single Nucleotide Variants

Microbial Single Nucleotide Variants (SNVs) were called using *metaSNV* (Costea et al., 2017). Each potential SNV required support by at least two non-reference sequencing reads (relative to the *specI* cluster representative genomes (Mende et al., 2013)) at a base call quality of Phred ≥20. The resulting sets of raw SNVs per taxon were filtered differentially for the various downstream analyses, as detailed below.

### Detection of Intra-Individual Microbial Transmission

To distinguish intra-individual microbial transmission from random drift, we calculated a *transmission score (S_T_)* per subject and microbial taxon. In short, S_T_ quantifies how much the similarity between oral and gut SNV profiles *within* an individual deviates from an *inter*-individual background. To calculate S_T_ we first filtered the set of informative SNVs (all SNVs at a given genome position) by applying the following criteria: (i) observation (read coverage ≥1) at focal position in ≥10 oral and ≥10 gut samples; (ii) SNV observation in ≥1 oral and ≥1 gut sample. Next, we calculated the global background incidence of each allele across oral (*f_oral_*) and gut (*f_gut_*) samples. From these, we calculated the background probabilities for each of the four possible cases in paired oral and gut observations: any given allele *i* could either be present in both samples (*p_1_,_1_*), absent in both samples (*p_0_,_0_*), or present in one but absent in the other sample (*p_1,0_* and *p_0,1_*):

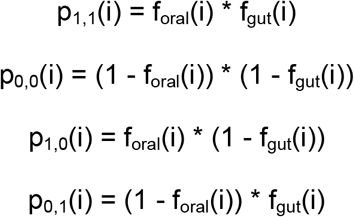

For every permuted oral-gut pair of samples, we then calculated the raw summed log-likelihood of the observed SNV profile overlap (*L_obs_*) across all alleles with shared coverage:

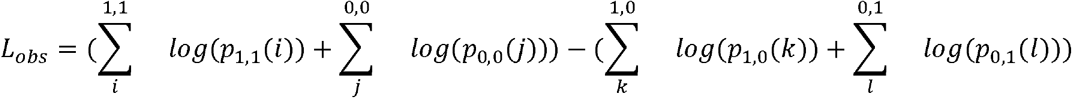

In other words, *L_obs_* quantifies how likely the observed average allele profile agreement between two samples is, given the respective background allele incidence frequencies. Similarly, we computed the log-likelihood of the least likely agreement case (*L_min_*) per allele:

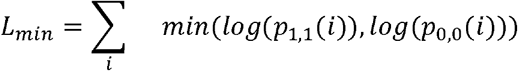

From these values, we calculated a raw probability score (*P_raw_*) for the observed allele agreement between a given pair of oral and gut samples:

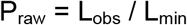

*P_raw_* scales the likelihood of the observed agreement by the likelihood of the theoretically most extreme cases of agreement across all observed alleles. In particular, the shared observation of very rare alleles (very low *f_oral_* and *f_gut_*) has a strong impact on *P_raw_*, whereas the shared observation of very common variants is downweighted.

We computed *P_raw_* for all pairwise permutations of oral and gut samples in the dataset with observations (reads) at ≥20 matching positions. We defined the transmission score *S_T_(t, s)* for taxon *t* in subject *s* as a standard Z score of the *intra*-individual (within subject) observation against an *inter*-individual (between subjects) background:

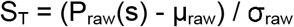

We tested for potential effects of the choice of background observations by calculating *ST* against (i) a global background of all pairwise inter-individual oral-gut comparisons, across all cohorts; (ii) a cohort-specific background per subject; (iii) a global background, but taking only subject-specific comparisons into account (the focal subject’s oral sample vs all gut samples, and vice versa); (iv) a within-cohort subject-specific background. Oral-gut comparisons for the same individual across different timepoints, within families (information available for LU and CN cohorts) and within village (for the Fijian cohort) were excluded from the background sets. Although smaller background sets (iii and iv) provided generally noisier scores, overall trends between these backgrounds were very consistent; in particular, cohort-specific vs global backgrounds did not impact trends in our findings (data not shown). All results discussed in the main text therefore refer to scores against a cohort-specific background (ii).

### Quantification of Intra-Individual Microbial Transmission

To quantify oral-gut transmission per individual, we defined a set of potentially *transmissible* species to include both frequently and occasionally transmitting species. Frequent transmitters encompassed a set of 74 species for which intra-individual transmission scores ST across subjects were significantly higher than inter-individual background (Benjamini-Hochberg-adjusted one-sided Wilcoxon p<0.05). Occasional transmitters did not satisfy this global criterion, but showed significant evidence for oral-fecal strain overlap in at least one individual (Benjamini-Hochberg-adjusted Z test p<0.05).

To quantify the transmitted microbial abundance per individual, we adjusted the observed relative oral and fecal abundance of each given species by oral-fecal SNV overlap. In other words, the *potentially* transmissible abundance in the oral cavity was defined as the total abundance of potentially transmitting species, and the *realised* transmitted abundance was defined to include only species for which overlapping strain populations could be confidently traced within individuals. This included frequent transmitters that were observable (above detection limits) in matched oral-fecal sample pairs, and occasional transmitters satisfying the additional criterion that significant transmission scores were required in the focal individual for (i.e., an occasional transmitter such as *Prevotella denticola* would only be considered in individuals in which it showed significant transmission scores). For these species, relative oral and fecal abundances were adjusted for total strain population overlap, estimated as the Jaccard overlap of SNVs observed in the oral cavity and gut of the focal individual.

### Longitudinal Coupling of Oral and Fecal SNV profiles

Longitudinal data (2-3 timepoints, see table S1) was available for 57 individuals from 3 cohorts (Heintz-Buschart et al., 2016; Voigt et al., 2015; Zhang et al., 2015). To quantify site-specific temporal stability of strain populations, we contrasted within-subject SNV profile similarity over time to between-subject similarities.

Moreover, we tested the longitudinal coupling of strain populations between a putative source site (e.g., oral cavity) and sink site (e.g., gut). For this, we required shared observations (read coverage ≥1) for at least 100 SNV positions across three samples (see Figure S1): (i) source site at the initial time point (t_0_); (ii) sink site at t_0_; (iii) sink site at a later time point t_1_. We defined source SNVs as present in sample (i), and newly gained sink SNVs as present in sample (iii) but not (ii), and performed Fisher’s exact tests (followed by Benjamini-Hochberg correction) to test for associations between these SNV sets. In other words, we tested for the association of strain populations present in the source site at t_0_ with strains newly gained in the sink site over time, by proxy of SNV profiles. We considered two sites to be longitudinally coupled in the source -> sink direction if the tested odds ratio was >1 at a (corrected) p≤0.05. Significant odds ratios <1 indicated unconnected sites in the tested directionality. Tests were performed independently for oral-to-gut (oral as source, gut as sink) and gut-to-oral coupling, per each taxon.

### Quantification of Oral-Fecal Transmission Rates

Longitudinal data was also leveraged to estimate oral-fecal transmission rates, here defined as the fraction of fecal strain turnover attributable to the corresponding salivary sample. For each subject and taxon, the absolute fecal strain turnover was quantified as described above, as the difference in SNV profiles between fecal samples at t_0_ and t_1_ (samples ii and iii in the previous section). Though sampling intervals ranged from 1 week to >1 year, they were relatively consistent within cohorts (see table S1). Transmission rates were then quantified as the fraction of fecal alleles gained between t_0_ and t_1_ that were also observed in the paired oral sample at t_0_. Arguably, this provides a conservative lower estimate: oral-fecal transmission could account for both newly gained fecal alleles and for the enhanced stability of existing alleles in the fecal strain population due to a constantly exerted dispersal pressure. However, since the latter effect cannot reliably be quantified from sparse longitudinal metagenomic data, the transmission rates reported in the main text only encompass the former (newly gained alleles).

To test whether transmission rates per taxon were statistically significant across subjects, we compared observed rates to two distinct randomised backgrounds: by shuffling fecal samples at t_1_ within cohorts, subject-specific *longitudinal* background sets on fecal strain turnover were generated; shuffling oral samples at t_0_ provided subject-specific *coupled* backgrounds. For each taxon and subject, we Z-transformed observed transmission rates against either of these subject-specific backgrounds; the resulting standard scores (in unit standard deviations) are reported in Figure 1C.

### Diversity, Community Composition and Statistical Analyses

Per-sample community richness was calculated from the average of 100 rarefactions to normalised marker gene-based abundances of 1,000. Between-sample community compositional similarities were computed as Bray-Curtis and TINA indices, as described previously (Schmidt et al., 2016). Distance-based Redundancy Analyses to associate community composition to levels of oral-fecal transmission were performed using the R package *vegan* (Oksanen et al., 2015).

The association of transmission scores with taxa phenotypes (oxygen requirement, sporulation, etc.) and taxa disease annotations (opportunistic pathogenicity, etc.) were tested using ANOVA of a combined linear model (‘naïve’ ANOVA in table S2). To correct for potentially confounding phylogenetic signals of the tested variables, an ANOVA of a phylogenetically regressed model of the same formulation was performed using the R package *caper* (Orme et al., 2018).

Associations of total transmitted classifiable abundance in saliva and stool per subject with subject variables (sex, BMI, age) were tested using ANOVAs on linear models blocked by cohort. The association of transmission scores per subject with disease status was tested using ANOVAs per disease cohort, on linear models accounting for taxon baselines, as well as effects of subject sex, BMI and age.

To test for links between microbiome composition and the amount of transmitted abundance in saliva and stool, we trained machine learning models to classify samples into ‘high’ and ‘low’ transmission groups. These groups were defined as the top and bottom quartiles of the fraction of transmitted abundance, independently for stool and saliva samples. For model training, relative abundances were log-transformed and standardized as z-scores. In a 10 times-repeated 10-fold cross-validation setting, L1-regularized (LASSO) logistic regression models (Tibshirani, 1996) were trained on the training set and then evaluated on the test set within each fold. In a second step, all species defined as frequent transmitters (see Quantification of Intra-Individual Microbial Transmission above) were eliminated as features before preprocessing and training. All steps (data preprocessing, model building, and model evaluation) were performed using the SIAMCAT R package (https://bioconductor.org/packages/SIAMCAT, version 1.1.0; see also Zeller et al, 2014).

All statistical analyses were performed in R. Analysis code is available online (see below).

### Data and Analysis Code Availability

All generated raw sequence data has been uploaded to the *European Nucleotide Archive* under the project accessions PRJEB28422 (French CRC, (Zeller et al., 2014) and German German healthy controls, (Voigt et al., 2015)) and PRJNA289586 (Luxembourg T1D, (Heintz-Buschart et al., 2016)). Sample metadata is available from table S1. Processed data (taxonomic profiles, taxa annotations, etc.) and full analysis code are available via a gitlab repository (https://git.embl.de/tschmidt/oral-fecal-transmission-public).

## Acknowledgements

The authors would like to thank Sina Klai of the University of Zürich, Switzerland, Johanna M. Schmidt and Gereon Rieke of the University of Bonn, Germany, for helpful comments and discussions on this manuscript, in particular regarding the medical relevance of several of the discussed bacterial species. We thank Katri Korpela, Lucas Silva, Thea van Rossum and other members of the Bork lab at EMBL, Germany, for helpful discussions. We thank Anna M Glazek and Yang Ping Yuan for bioinformatics support, Stefanie Kandels-Lewis of the EMBL for support on sample logistics and administration, Rajna Hercog, Jan Provaznik and Vladimir Benes and, in general, the EMBL Genomics Core Facility for sequencing support, and Laura Lebrun of LCSB for support with the biomolecular extraction platform. TSBS, MRH and AHB were supported by a Luxembourg National Research Fund CORE-INTER grant (MicroCancer; CORE/15/BM/10404093). MRH was additionally supported by a Marie Curie Individual Fellowship (661019). TSBS, SSL, OMM, RJA and PB were supported by an European Research Council grant (MicroBioS; ERC-AdG-669830). GZ and PB were supported by the BMBF-funded Heidelberg Center for Human Bioinformatics (HD-HuB) within the German Network for Bioinformatics Infrastructure (de.NBI #031A537B).

## References

Asnicar F, Manara S, Zolfo M, Truong DT, Scholz M, Armanini F, Ferretti P, Gorfer V, Pedrotti A, Tett A, Segata N, Gilbert JA. 2017. Studying Vertical Microbiome Transmission from Mothers to Infants by Strain-Level Metagenomic Profiling. mSystems 2:e00164–16.

Atarashi K, Suda W, Luo C, Kawaguchi T, Motoo I, Narushima S, Kiguchi Y, Yasuma K, Watanabe E, Tanoue T, Thaiss CA, Sato M, Toyooka K, Said HS, Yamagami H, Rice SA, Gevers D, Johnson RC, Segre JA, Chen K, Kolls JK, Elinav E, Morita H, Xavier RJ, Hattori M, Honda K. 2017. Ectopic colonization of oral bacteria in the intestine drives TH1 cell induction and inflammation. Science 358:359–365.

Brito IL, Yilmaz S, Huang K, Xu L, Jupiter SD, Jenkins AP, Naisilisili W, Tamminen M, Smillie CS, Wortman JR, Birren BW, Xavier RJ, Blainey PC, Singh AK, Gevers D, Alm EJ. 2016. Mobile genes in the human microbiome are structured from global to individual scales. Nature 535:435–439.

Coelho LP, Alves R, Monteiro P, Huerta-Cepas J, Freitas AT, Bork P. 2018. NG-meta-profiler: fast processing of metagenomes using NGLess, a domain-specific language. bioRxiv. doi:10.1101/367755

Costea PI, Munch R, Coelho LP, Paoli L, Sunagawa S, Bork P. 2017. metaSNV: A tool for metagenomic strain level analysis. PLoS One 12:e0182392.

Ding T, Schloss PD. 2014. Dynamics and associations of microbial community types across the human body. Nature 509:357–360.

Felmer B, Warren RD, Barrett MP, Cisek K, Das A, Jeffery IB, Hurley E, O’Riordain M, Shanahan F, O’Toole PW. 2018. The oral microbiota is distinctive and predictive. Gut 67:1454–1463.

Ferretti P, Pasolli E, Tett A, Asnicar F, Gorfer V, Fedi S, Armanini F, Truong DT, Manara S, Zolfo M, Beghini F, Bertorelli R, De Sanctis V, Bariletti I, Canto R, Clementi R, Cologna M, Crifò T, Cusumano G, Gottardi S, Innamorati C, Masè C, Postai D, Savoi D, Duranti S, Lugli GA, Mancabelli L, Turroni F, Ferrario C, Milani C, Mangifesta M, Anzalone R, Viappiani A, Yassour M, Vlamakis H, Xavier R, Collado CM, Koren O, Tateo S, Soffiati M, Pedrotti A, Ventura M, Huttenhower C, Bork P, Segata N. 2018. Mother-to-Infant Microbial Transmission from Different Body Sites Shapes the Developing Infant Gut Microbiome. Cell Host Microbe 24:133–145.e5.

Flynn KJ, Baxter NT, Schloss PD, McMahon K. 2016. Metabolic and Community Synergy of Oral Bacteria in Colorectal Cancer 1:e00102–16.

Franzosa EA, Morgan XC, Segata N, Waldron L, Reyes J, Earl AM, Giannoukos G, Boylan MR, Ciulla D, Gevers D, Izard J, Garrett WS, Chan AT, Huttenhower C. 2014. Relating the metatranscriptome and metagenome of the human gut. Proc Natl Acad Sci U S A 111:E2329–38.

Gevers D, Kugathasan S, Denson LA, Vázquez-Baeza Y, Van Treuren W, Ren B, Schwager E, Knights D, Song SJ, Yassour M, Morgan XC, Kostic AD, Luo C, Gonzalez A, McDonald D, Haberman Y, Walters T, Baker S, Rosh J, Stephens M, Heyman M, Markowitz J, Baldassano R, Griffiths A, Sylvester F, Mack D, Kim S, Crandall W, Hyams J, Huttenhower C, Knight R, Xavier RJ. 2014. The Treatment-Naive Microbiome in New-Onset Crohn’s Disease. Cell Host Microbe 15:382–392.

Giannella RA, Broitman SA, Zamcheck N. 1972. Gastric acid barrier to ingested microorganisms in man: studies in vivo and in vitro. Gut 13:251–256.

Hajishengallis G. 2014. Periodontitis: from microbial immune subversion to systemic inflammation. Nat Rev Immunol 15:30–44.

Heintz-Buschart A, May P, Laczny CC, Lebrun LA, Bellora C, Krishna A, Wampach L, Schneider JG, Hogan A, de Beaufort C, Wilmes P. 2016. Integrated multi-omics of the human gut microbiome in a case study of familial type 1 diabetes. Nature Microbiology 2:16180.

Howden CW, Hunt RH. 1987. Relationship between gastric secretion and infection. Gut 28:96–107.

Huerta-Cepas J, Serra F, Bork P. 2016. ETE 3: Reconstruction, Analysis, and Visualization of Phylogenomic Data. Mol Biol Evol 33:1635–1638.

Humphrey SP, Williamson RT. 2001. A review of saliva: normal composition, flow, and function. J Prosthet Dent 85:162–169.

Imhann F, Bonder MJ, Vila AV, Fu J, Mujagic Z, Vork L, Tigchelaar EF, Jankipersadsing SA, Cenit MC, Harmsen HJM, Dijkstra G, Franke L, Xavier RJ, Jonkers D, Wijmenga C, Weersma RK, Zhernakova A. 2016. Proton pump inhibitors affect the gut microbiome. Gut 65:740–748.

Jones DT, Taylor WR, Thornton JM. 1992. The rapid generation of mutation data matrices from protein sequences. Comput Appl Biosci 8:275–282.

Korpela K, Costea PI, Coelho LP, Kandels-Lewis S, Willemsen G, Boomsma DI, Segata N, Bork P. 2018. Selective maternal seeding and environment shape the human gut microbiome. Genome Res. doi:10.1101/gr.233940.117

Kultima JR, Coelho LP, Forslund K, Huerta-Cepas J, Li SS, Driessen M, Voigt AY, Zeller G, Sunagawa S, Bork P. 2016. MOCAT2: a metagenomic assembly, annotation and profiling framework. Bioinformatics 32:2520–2523.

Letunic I, Bork P. 2016. Interactive tree of life (iTOL) v3: an online tool for the display and annotation of phylogenetic and other trees. Nucleic Acids Res 44:W242–5.

Li H, Durbin R. 2009. Fast and accurate short read alignment with Burrows–Wheeler transform. Bioinformatics 25:1754–1760.

Li SS, Zhu A, Benes V, Costea PI, Hercog R, Hildebrand F, Huerta-Cepas J, Nieuwdorp M, Salojärvi J, Voigt AY, Zeller G, Sunagawa S, de Vos WM, Bork P. 2016. Durable coexistence of donor and recipient strains after fecal microbiota transplantation. Science 352:586–589.

Liu Y, Zhang Y, Dong P, An R, Xue C, Ge Y, Wei L, Liang X. 2015. Digestion of Nucleic Acids starts in the Stomach. Scientific Reports 5:11936

Lloyd-Price J, Mahurkar A, Rahnavard G, Crabtree J, Orvis J, Hall AB, Brady A, Creasy HH, McCracken C, Giglio MG, McDonald D, Franzosa EA, Knight R, White O, Huttenhower C. 2017. Strains, functions and dynamics in the expanded Human Microbiome Project. Nature 486:207.

Lynch SV, Pedersen O. 2016. The Human Intestinal Microbiome in Health and Disease. N Engl J Med 375:2369–2379.

Martinsen TC, Bergh K, Waldum HL. 2005. Gastric juice: a barrier against infectious diseases. Basic Clin Pharmacol Toxicol 96:94–102.

Mende DR, Sunagawa S, Zeller G, Bork P. 2013. Accurate and universal delineation of prokaryotic species. Nature Publishing Group 10:881–884.

Mercer DK, Scott KP, Bruce-Johnson WA, Glover LA, Flint HJ. 1999. Fate of Free DNA and Transformation of the Oral Bacterium *Streptococcus gordonii* DL1 by Plasmid DNA in Human Saliva. Appl Environ Microbiol 65(1):6–10

Oksanen J, Blanchet FG, Kindt R, Legendre P, Minchin PR, O’Hara RB, Simpson GL, Solymos P, Stevens MHS, Wagner H. 2015. vegan: Community Ecology Package.

Orme D, Freckleton R, Thomas G, Petzoldt T, Fritz S, Isaac N, Pearse W. 2018. caper: Comparative Analyses of Phylogenetics and Evolution in R.

Price MN, Dehal PS, Arkin AP. 2010. FastTree 2 – Approximately Maximum-Likelihood Trees for Large Alignments. PLoS One 5:e9490.

Ridlon JM, Kang DJ, Hylemon PB, Bajaj JS. 2014. Bile acids and the gut microbiome. Curr Opin Gastroenterol 30:332–338.

Savage DC. 1977. Microbial Ecology of the Gastrointestinal Tract. Annu Rev Microbiol 31:107–133.

Schloissnig S, Arumugam M, Sunagawa S, Mitreva M, Tap J, Zhu A, Waller A, Mende DR, Kultima JR, Martin J, Kota K, Sunyaev SR, Weinstock GM, Bork P. 2012. Genomic variation landscape of the human gut microbiome. Nature 493:45–50.

Schmidt TSB, Raes J, Bork P. 2018. The Human Gut Microbiome: From Association to Modulation. Cell 172:1198–1215.

Schmidt TSB, Rodrigues JFM, von Mering C. 2016. A family of interaction-adjusted indices of community similarity. ISME J.

Segata N, Haake SK, Mannon P, Lemon KP, Waldron L, Gevers D, Huttenhower C, Izard J. 2012. Composition of the adult digestive tract bacterial microbiome based on seven mouth surfaces, tonsils, throat and stool samples. Genome Biol 13:R42.

Sender R, Fuchs S, Milo R. 2016. Revised Estimates for the Number of Human and Bacteria Cells in the Body. PLoS Biol 14:e1002533.

Sievers F, Wilm A, Dineen D, Gibson TJ, Karplus K, Li W, Lopez R, McWilliam H, Remmert M, Söding J, Thompson JD, Higgins DG. 2011. Fast, scalable generation of high-quality protein multiple sequence alignments using Clustal Omega. Mol Syst Biol 7:539.

Socransky SS, Haffajee AD, Cugini MA, Smith C, Kent RL. 1998. Microbial complexes in subgingival plaque. J Clin Periodontol 25:134–144.

Tibshirani R., Regression Shrinkage and Selection via the Lasso. J.R. Stat. Soc. Series B Stat. Methodol., 1996. 58(1): p. 267–288.

Voigt AY, Costea PI, Kultima JR, Li SS, Zeller G, Sunagawa S, Bork P. 2015. Temporal and technical variability of human gut metagenomes. Genome Biol 16:1.

Wade WG. 2013. The oral microbiome in health and disease. Pharmacol Res 69:137–143.

Wattam AR, Davis JJ, Assaf R, Boisvert S, Brettin T, Bun C, Conrad N, Dietrich EM, Disz T, Gabbard JL, Gerdes S, Henry CS, Kenyon RW, Machi D, Mao C, Nordberg EK, Olsen GJ, Murphy-Olson DE, Olson R, Overbeek R, Parrello B, Pusch GD, Shukla M, Vonstein V, Warren A, Xia F, Yoo H, Stevens RL. 2017. Improvements to PATRIC, the all-bacterial Bioinformatics Database and Analysis Resource Center. Nucleic Acids Res 45:D535–D542.

Zeller G, Tap J, Voigt AY, Sunagawa S, Kultima JR, Costea PI, Amiot A, Böhm J, Brunetti F, Habermann N, Hercog R, Koch M, Luciani A, Mende DR, Schneider MA, Schrotz King P, Tournigand C, Tran Van Nhieu J, Yamada T, Zimmermann J, Benes V, Kloor M, Ulrich CM, Knebel Doeberitz M, Sobhani I, Bork P. 2014. Potential of fecal microbiota for early-stage detection of colorectal cancer. Mol Syst Biol 10:766–766.

Zhang X, Zhang D, Jia H, Feng Q, Wang D, Di Liang, Wu X, Li J, Tang L, Li Y, Lan Z, Chen B, Li Y, Zhong H, Xie H, Jie Z, Chen W, Tang S, Xu X, Wang X, Cai X, Liu S, Xia Y, Li J, Qiao X, Al-Aama JY, Chen H, Wang L, Wu Q-J, Zhang F, Zheng W, Li Y, Zhang M, Luo G, Xue W, Xiao L, Li J, Chen W, Xu X, Yin Y, Yang H, Wang J, Kristiansen K, Liu L, Li T, Huang Q, Li Y, Wang J. 2015. The oral and gut microbiomes are perturbed in rheumatoid arthritis and partly normalized after treatment. Nat Med 21:895–905.

Zmora N, Zilberman-Schapira G, Suez J, Mor U, Dori-Bachash M, Bashiardes S, Kotler E, Zur M, Regev-Lehavi D, Brik RB, Federici S, Cohen Y, Linevsky R, Rothschild D, Moor AE, Ben-Moshe S, Harmelin A, Itzkovitz S, Maharshak N, Shibolet O, Shapiro H, Pevsner-Fischer M, Sharon I, Halpern Z, Segal E, Elinav E. 2018. Personalized Gut Mucosal Colonization Resistance to Empiric Probiotics Is Associated with Unique Host and Microbiome Features. Cell 174:1388–1405.

