## Supplementary figures and images for "“Extensive Transmission of Microbes along the Gastrointestinal Tract”"

### Supporting Figure S1

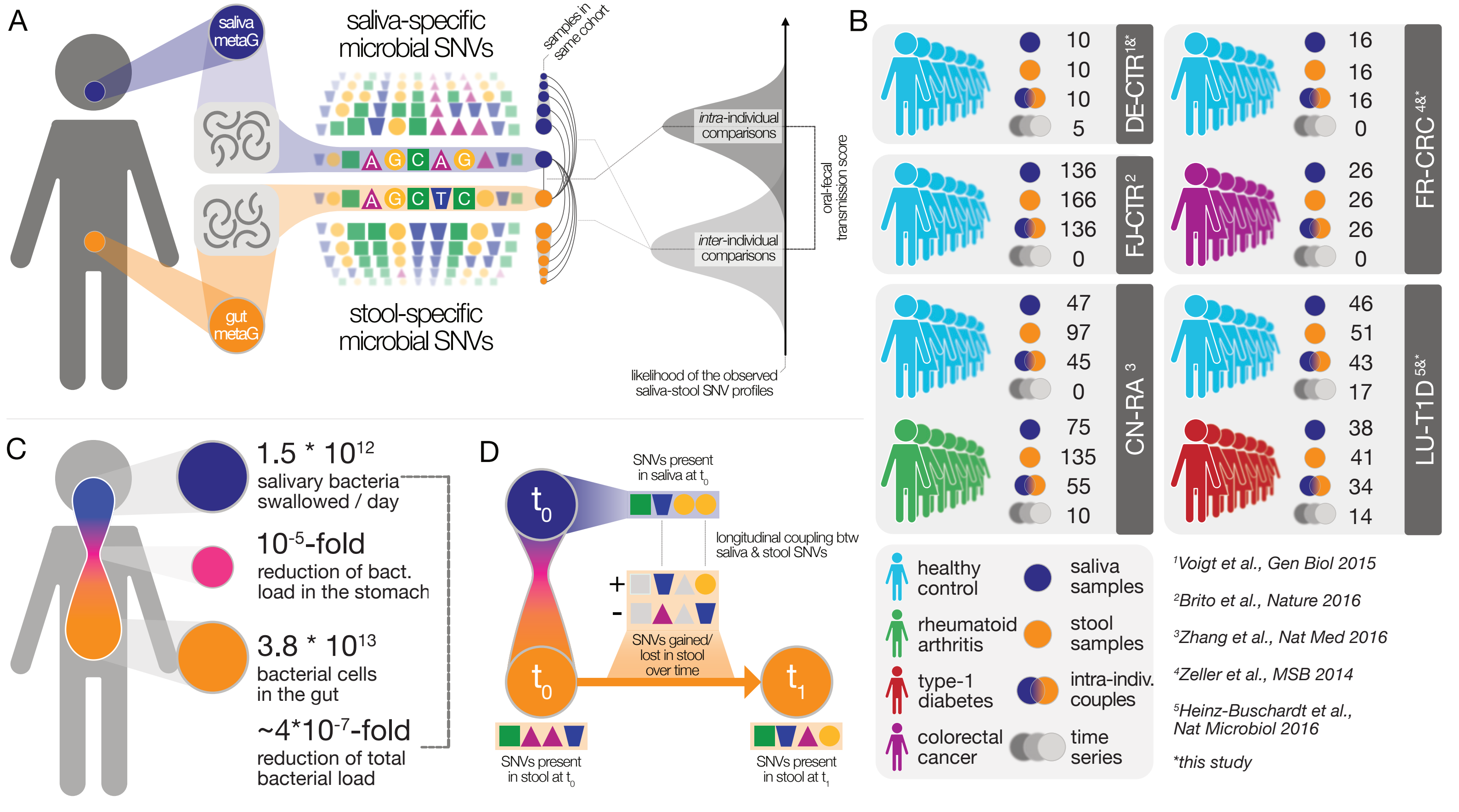

### Supporting Figure S2

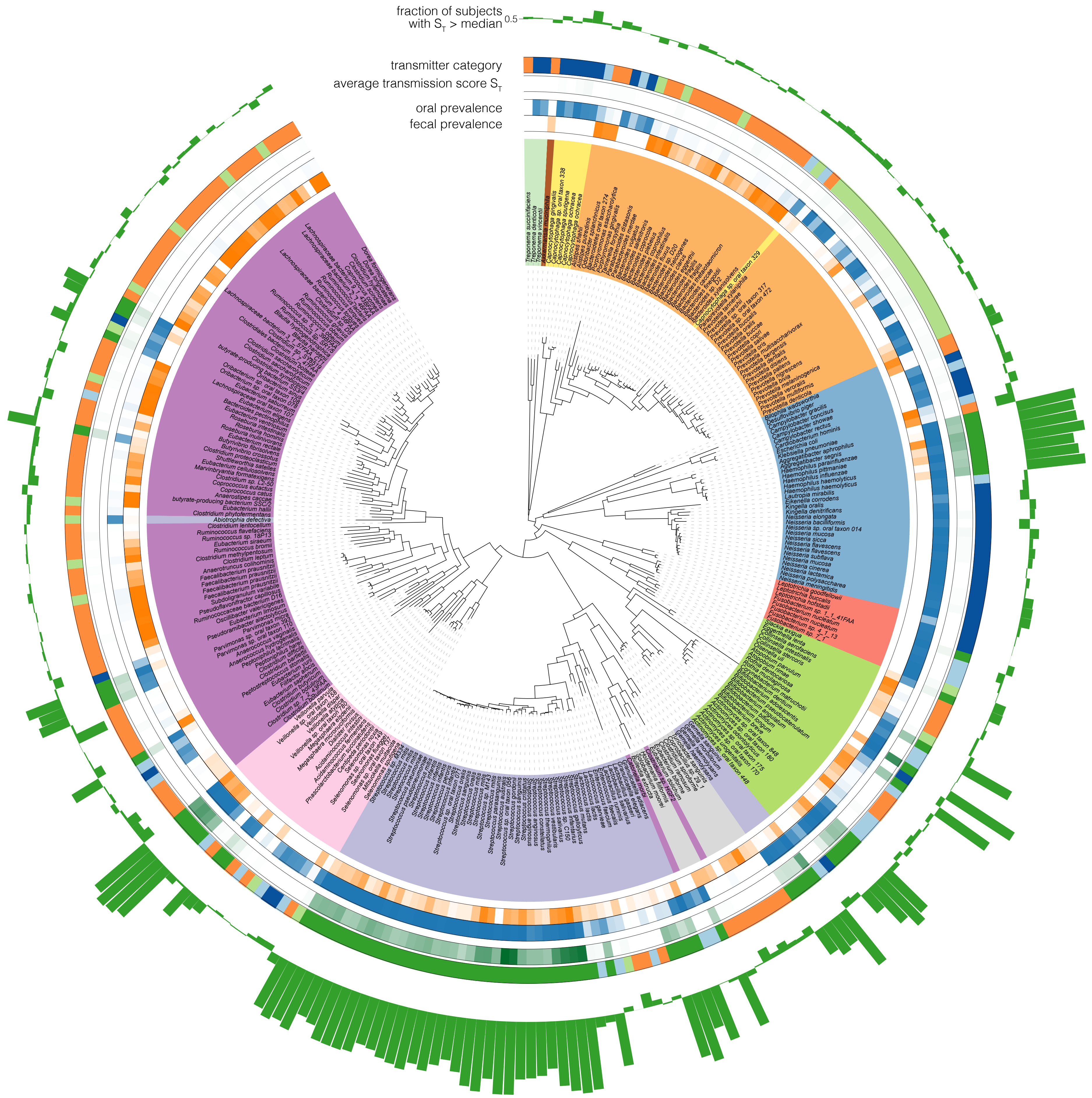

### Supporting Figure S3

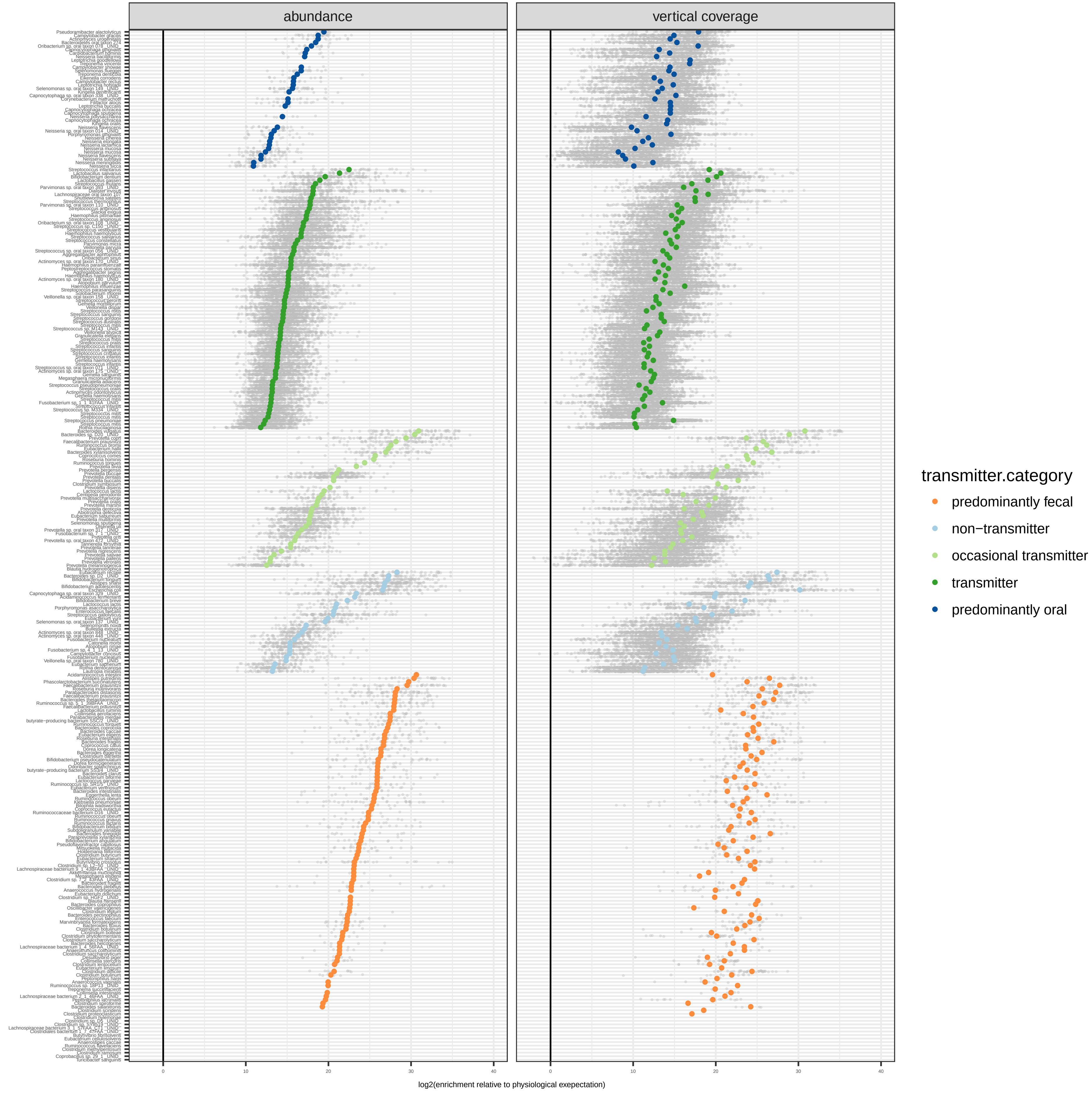

### Supporting Figure S4

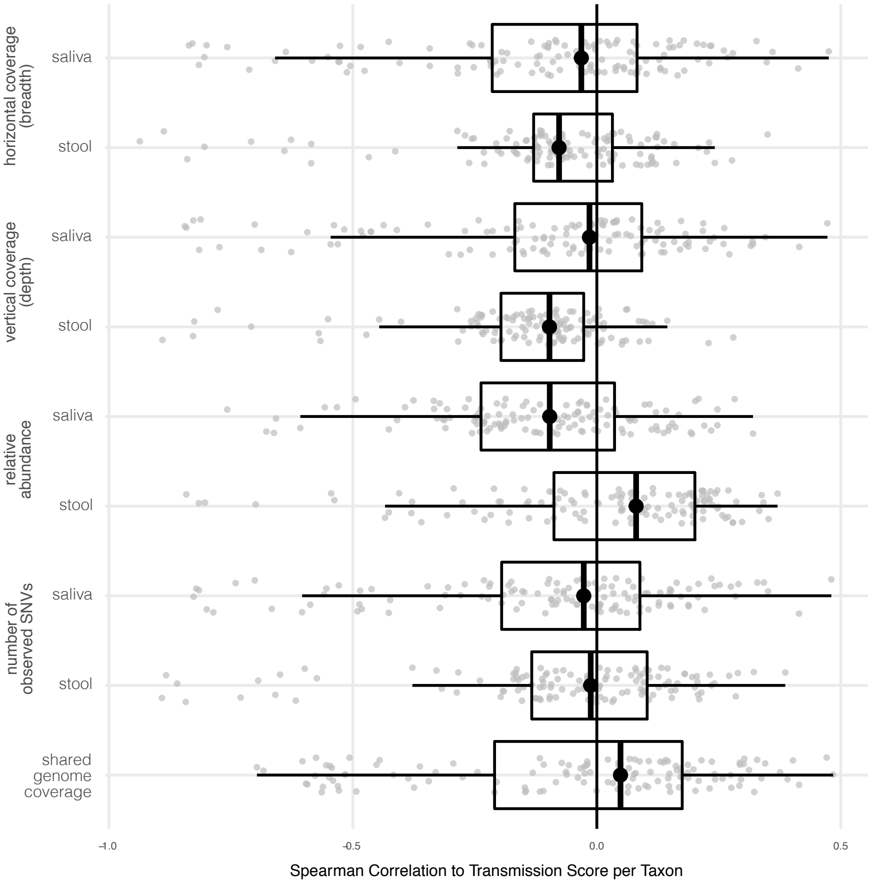

### Supporting Figure S5

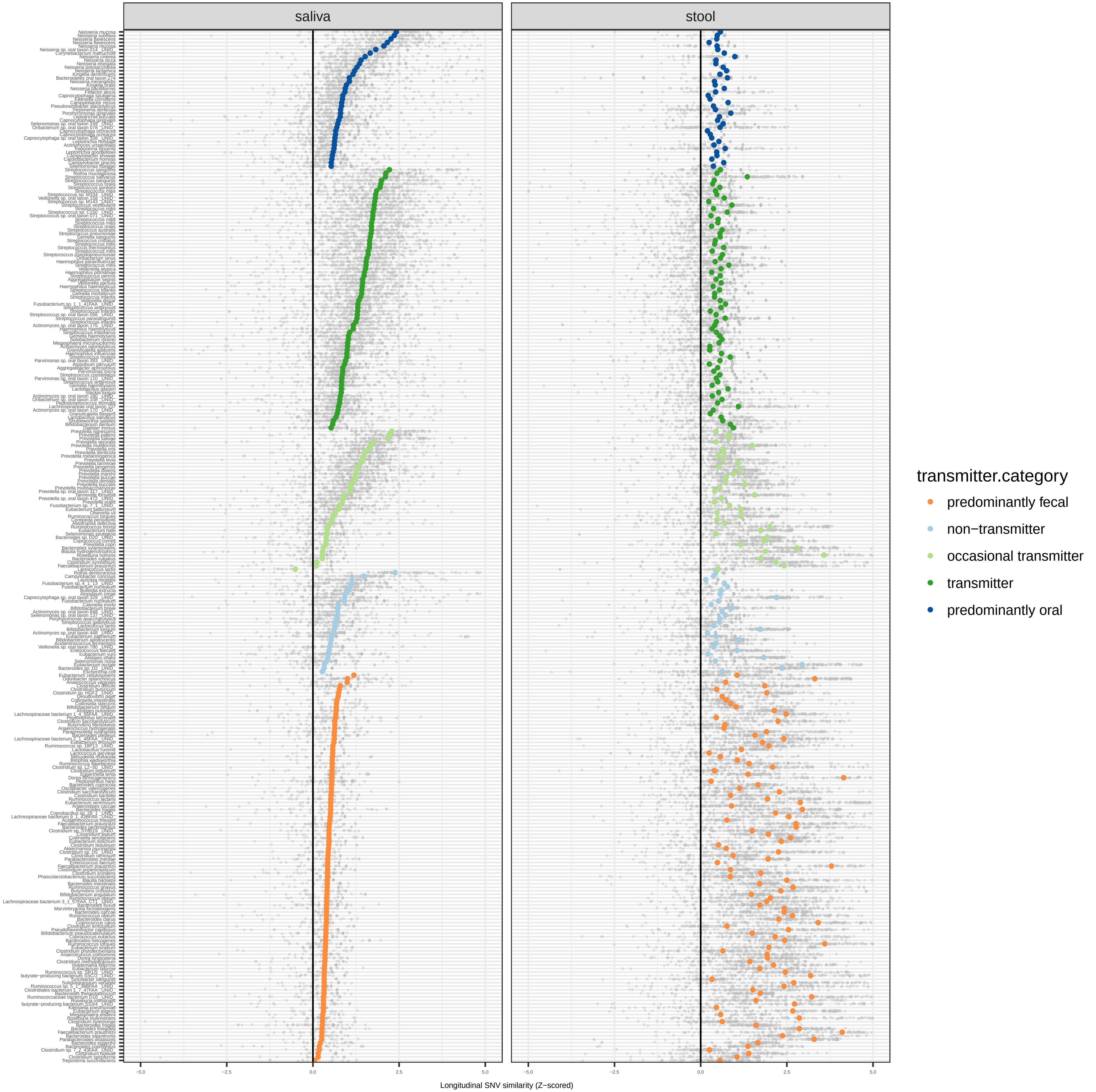

### Supporting Figure S6

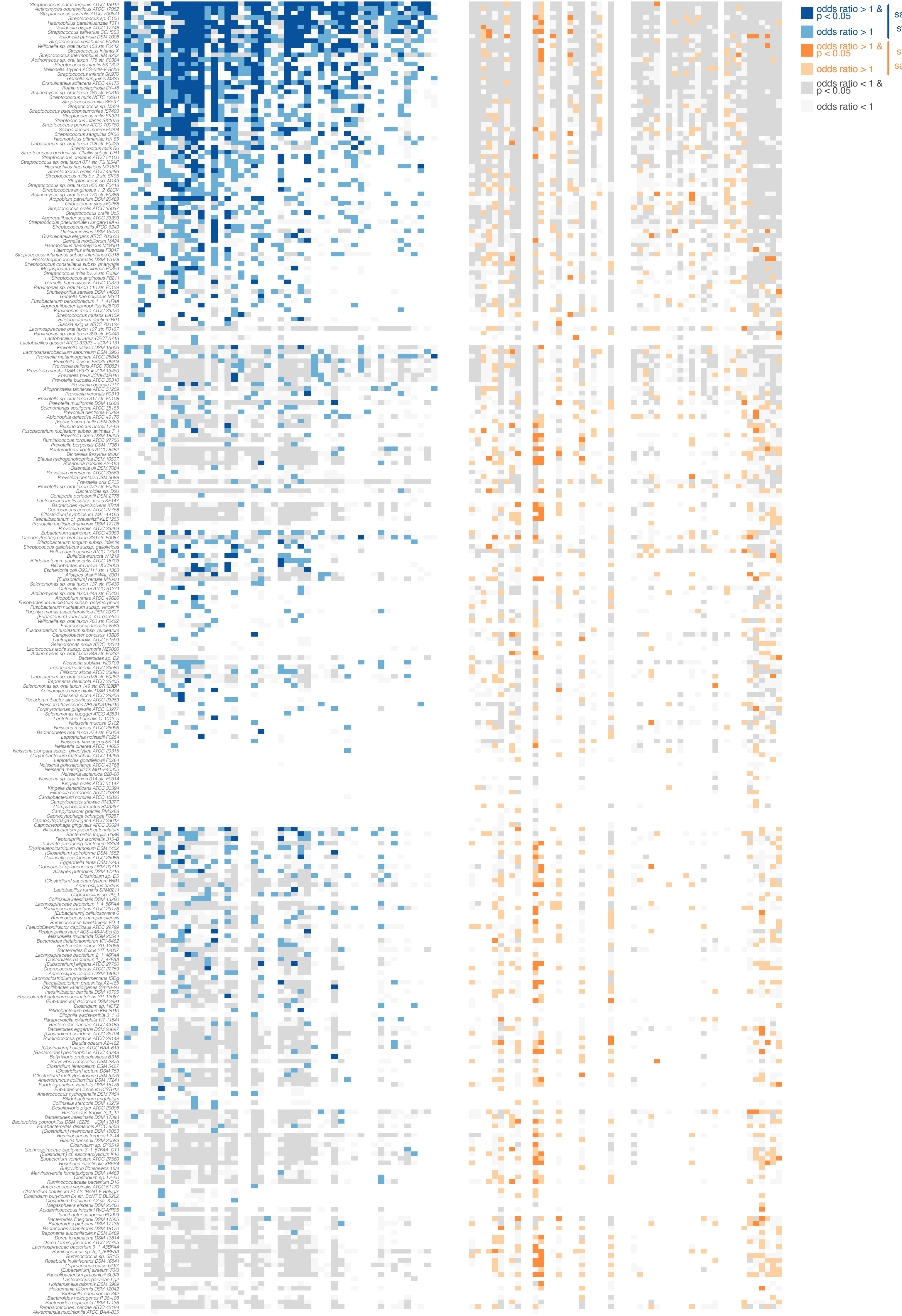

### Supporting Figure S7

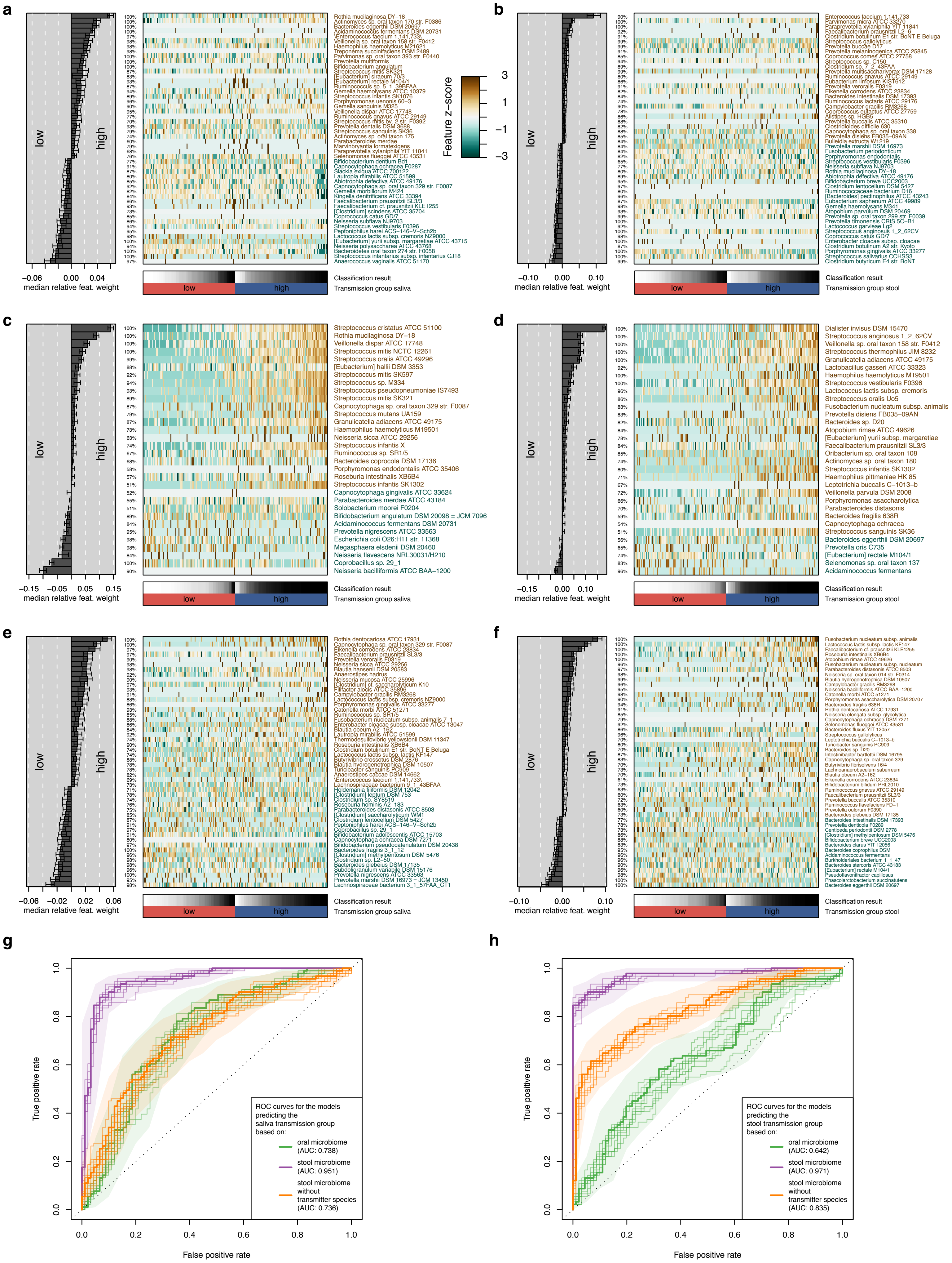

### Supporting Figure S8

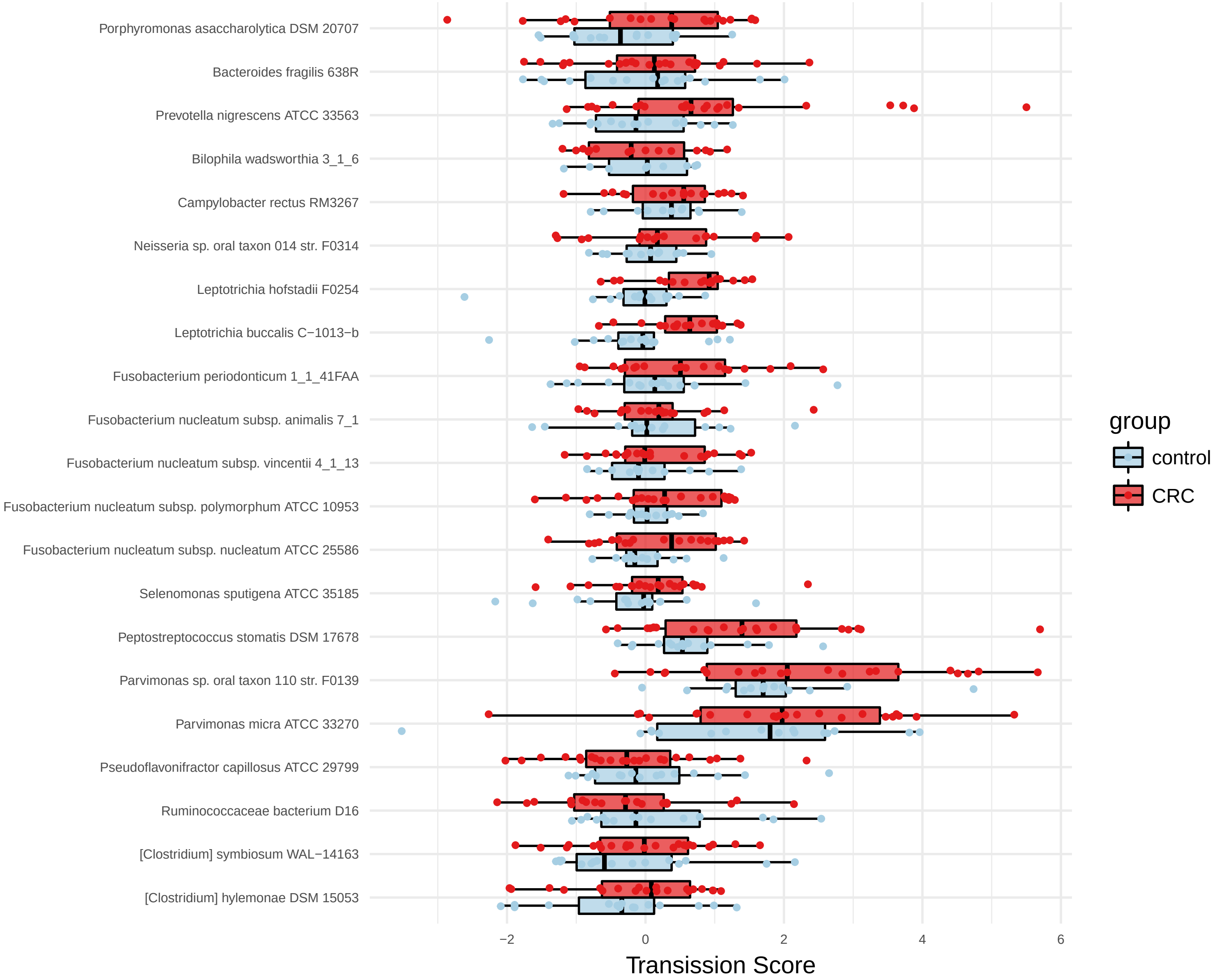

### Supporting Figure S9

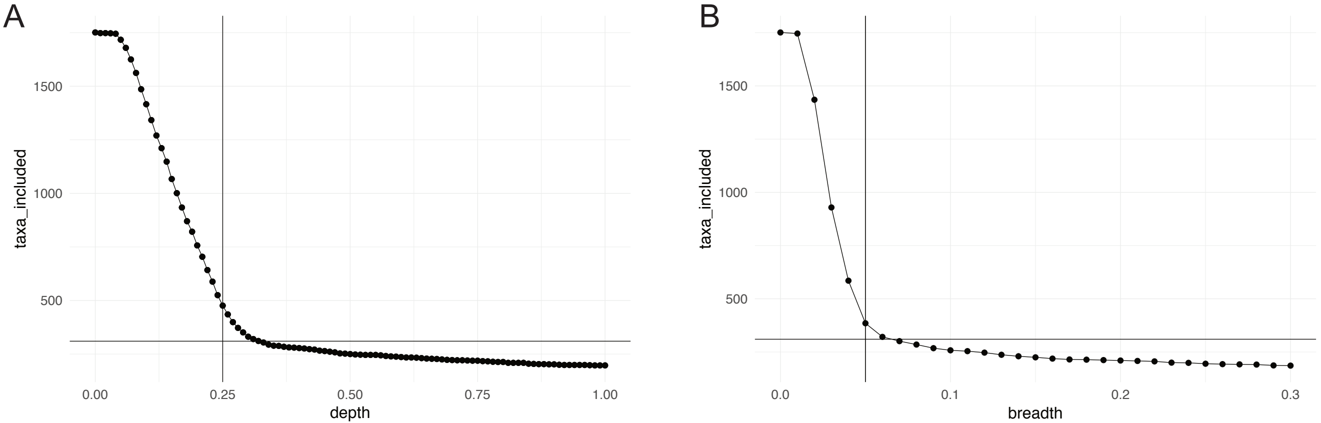
